# Decomposing spatially dependent and cell type specific contributions to cellular heterogeneity

**DOI:** 10.1101/275156

**Authors:** Qian Zhu, Sheel Shah, Ruben Dries, Long Cai, Guo-Cheng Yuan

## Abstract

Both the intrinsic regulatory network and spatial environment are contributors of cellular identity and result in cell state variations. However, their individual contributions remain poorly understood. Here we present a systematic approach to integrate both sequencing-and imaging-based single-cell transcriptomic profiles, thereby combining whole-transcriptomic and spatial information from these assays. We applied this approach to dissect the cell-type and spatial domain associated heterogeneity within the mouse visual cortex region. Our analysis identified distinct spatially associated signatures within glutamatergic and astrocyte cell compartments, indicating strong interactions between cells and their spatial environment. Using these signatures as a guide to analyze single cell RNAseq data, we identified previously unknown, but spatially associated subpopulations. As such, our integrated approach provides a powerful tool for dissecting the roles of intrinsic regulatory networks and spatial environment in the maintenance of cellular states.

## Introduction

Human and other multicellular organisms are composed of diverse cell types characterized by distinct gene expression patterns. Within each cell type, there is also considerable heterogeneity. The source of cellular heterogeneity remains poorly understood, but it is commonly thought to be modulated by the balance between intrinsic regulatory networks and extrinsic cellular microenvironment^1–5^. Recently, the rapid development of single-cell technologies has enabled accurate and simultaneous measurements of cell position and gene expression^6–9^, thus providing an excellent opportunity to systematically characterize cellular heterogeneity. However, the relative contribution of intrinsic and extrinsic factors in mediating cell-state variation remains poorly understood.

Currently, there are two major, complementary approaches for single-cell transcriptomic profiling. The first is single-cell RNA sequencing (scRNAseq)^6, 8, 10–15^. By combining single-cell isolation, library amplification, and massively parallel sequencing, scRNAseq provides the most comprehensive view of transcriptomes. The second approach is single-molecule fluorescence in situ hybridization (smFISH)^7, 16–20^, which can be used to detect mRNA transcripts with high sensitivity while maintaining the spatial content. With sequential rounds of smFISH imaging, it is now feasible to profile the expression level of hundreds of genes for each cell in tissues. Each technology features a distinct set of advantages and limitations. The sequential smFISH technology carries the advantage of measuring the transcriptome with high accuracy in its native spatial environment, but current implementations profile only a few hundred genes, whereas scRNAseq provides whole-transcriptome estimation but requires cells to be removed from their spatial environment, resulting in a loss of spatial information^19, 21^.

It is clear that an integrative analysis framework, involving scRNAseq and sequential smFISH, would bring together the benefits of both technologies to better characterize both cell type and spatially dependent variations. To this end, we developed a computational approach that contains two major components: First, the scRNAseq data is used as a guide to accurately determine the cell-types corresponding to the cells profiled by sequential smFISH. Second, distinct spatial domain patterns are systematically detected from sequential smFISH data. These spatial patterns are then in turn used to dissect the environment-associated variation in a scRNAseq dataset.

This integrated approach has enabled us to systematically dissect the respective contribution of cell type and spatially dependent factors in mediating cell-state variation (**Fig. 1a**), which has eluded previous studies. Most existing studies focused on identifying cell-type differences, but, as shown below in our analysis of the mouse visual cortex region, cell-type differences represent only one component in cell-state variation (schematically represented as the cell intrinsic dimension in **Fig. 1a**), whereas the spatial environment plays a significant role in mediating gene activities, probably through cell-cell interactions (represented as the spatial dimension in **Fig. 1a**) and signaling. As each technology has its own strengths and weaknesses, the integrated approach presented here provides a powerful model framework and broadly applicable to analyze diverse tissues from various model systems.

**Figure 1:**
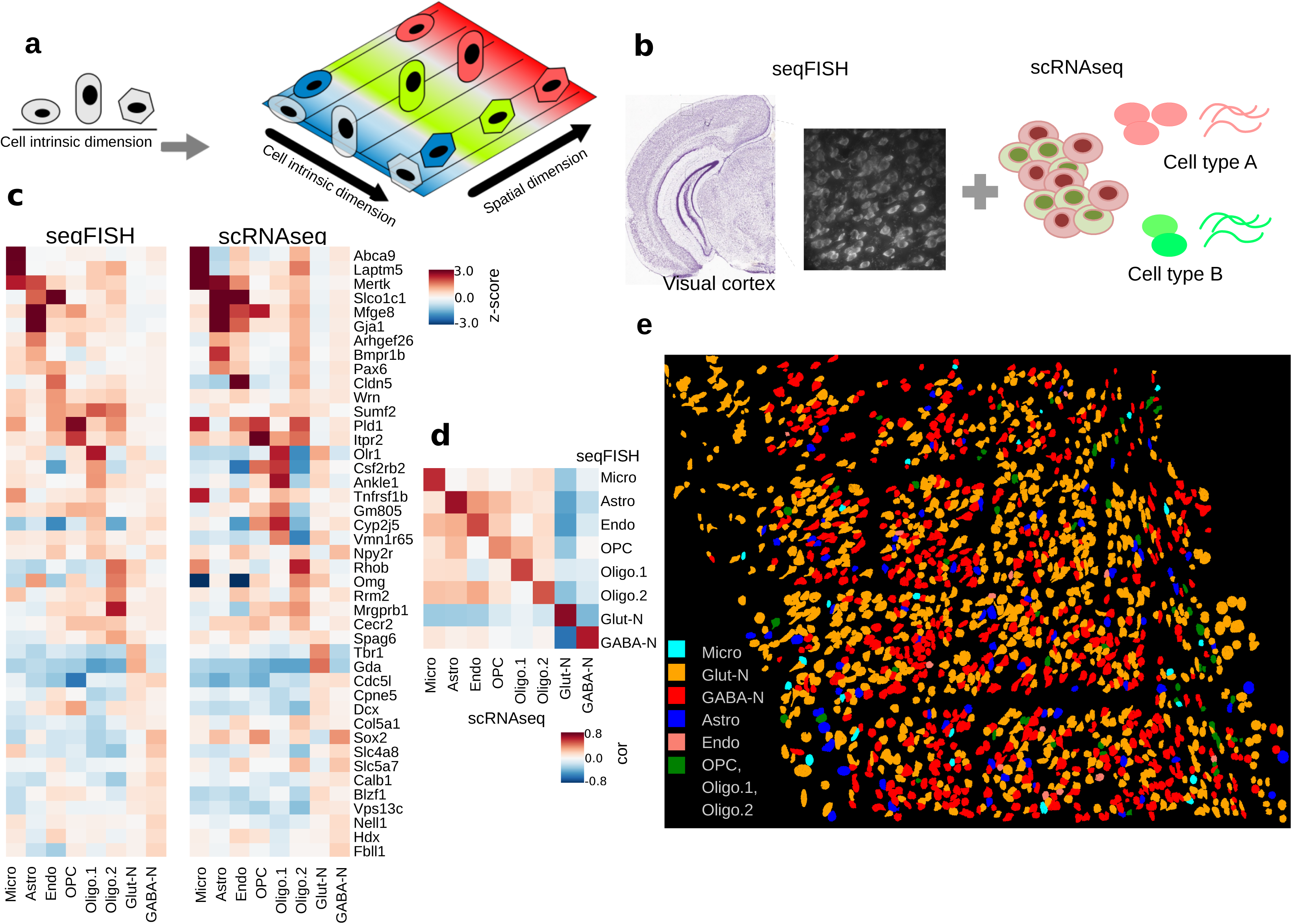
Overall goal of the project and cell type prediction in seqFISH data. a. Cellular heterogeneity is driven by both cell-type (indicated by shape) and environmental factors (indicated by colors). ScRNAseq based studies can only detect cell-type related variation, because spatial information is lost. b. Our goal is to decompose the contributions of each factor by developing methods to integrate scRNAseq and seqFISH data. c. Prediction results evaluated by the comparison of cell-type average expression profile across technologies for 8 major cell types. Values represent expression z-scores. d. Correlation between reference and predicted cell type averages ranges from 0.75 to 0.95. e. Integration of seqFISH and scRNAseq data (illustrated by b) enables cell-type mapping with spatial information in the adult mouse visual cortex. Each cell type is labeled by a different color. Cell shape information is obtained from segmentation of cells from images (see Methods).

## Results

### Mapping scRNAseq cell-types on seqFISH data

Given that scRNAseq, as a whole transcriptomic approach, can provide signatures for a diverse set of cell types, we took advantage of the whole-transcriptomic information obtained from scRNAseq data and developed a supervised cell-type mapping approach by integrating seqFISH and scRNAseq data (**Fig. 1b**). Our goal differs from previous studies^22–26^, where scRNAseq data were mapped onto conventional ISH images to predict cell locations. Of note, ISH images are not quantitative, multiplexed or single-cell resolution. In a seqFISH experiment, transcripts from hundreds of genes are detected directly in individual cells in their native spatial environment at single molecule resolution.

Our strategy is to use scRNAseq data to capture the large cell type differences and then further investigate spatial patterning within each major cell types. We analyzed a published scRNAseq dataset targeting the mouse visual cortex regions^27^. Eight major cell types: GABAergic, glutamatergic, astrocytes, 3 oligodendrocyte groups, microglia, and endothelial cells were identified from scRNAseq analysis^27^. To estimate the minimal number of genes that is required for accurate cell-type mapping, we randomly selected a subset from the list of differentially expressed (DE) genes across these cell types, and applied a multiclass support vector machine (SVM)^28, 29^ model using only the expression levels of these genes. The performance was evaluated by cross-validation. By using only 40 genes, we can already achieve an average level of 89% mapping accuracy. Not surprisingly, increasing the number of genes leads to better performance (92% for 60 genes, and 96% for 80 genes). Therefore, there is significant redundancy in transcriptomic profiles which can be compressed into fewer than 100 genes. We then investigated a seqFISH dataset for the mouse visual cortex area^19^. A 1 mm by 1 mm contiguous area of the mouse visual cortex was imaged with 4 barcoded rounds of hybridization to decode 100 unique transcripts followed by 5 rounds of non-combinatorial hybridization to quantify 25 highly expressed genes (**Supplementary Table 1**). These rounds of imaging were preceded by imaging of the DAPI stain in the region and followed by imaging of the Nissl stain in the region. The images were aligned and transcripts decoded as described in Shah et al. 2016. Transcripts were assigned to cells which were segmented based on Nissl and DAPI staining. Using this technology, we were able to quantify the expression levels of these 125 genes with high accuracy in a total of 1597 cells.

After computing differentially expressed genes across the 8 major cell types in Tasic et al, we selected the top 43 (P<1e-20) of these 125 genes for cell-type classification. These genes contain both highly expressed (>50 copies per cell) and lowly expressed genes (<10 copies per cell). Cross-validation analysis shows that, using these 43 genes as input, the SVM model accurately mapped 90.1% of the cells in the scRNAseq data to the correct cell-type. Therefore, we proceeded by using these 43 genes (**Supplementary Table 2**) to map cell-types in the seqFISH data.

As a first step, we preprocessed the seqFISH data by using a multi-image regression algorithm in order to reduce potential technical biases due to non-uniform imaging intensity variation (Methods). We further adopted a quantile normalization^30^ approach to calibrate the scaling and distribution differences between scRNAseq and seqFISH experiments. For most genes, the quantile-quantile (q-q) plot normalization curve is strikingly linear (**Supplementary Fig. 1**), suggesting a high degree of agreement between the two datasets despite technological differences. Then, the SVM classification model was applied to the bias-corrected, quantile-normalized seqFISH data to assign cell types. Of note, we found that better performance may be achieved by further calibrating model parameters to accommodate platform differences. The results of multiclass SVM are calibrated across models^31^ and converted to probabilities. The results showed the exclusion of 5.5% cells that cannot be confidently mapped to a single cell-type (with 0.5 or less probability). Among the mapped cells, 54% are glutamatergic neurons, 37% are GABAergic neurons, 4.8% are astrocytes, and other glial cell types and endothelial cells make up the remaining 4.2% of cells (**Fig. 1c**).

To validate our predictions, we first checked the expression of known marker genes and compared the average gene expression profiles between scRNAseq and seqFISH data. Indeed, this comparison shows a high degree of similarity (**Fig. 1c**). Notably, marker genes have expected high expression in the matched cell types, such as Gja1 and Mfge8 in astrocytes, Laptm5 and Abca9 in microglia, Cldn5 in endothelial cells, Tbr1 and Gda in glutamatergic neurons, and Slc5a7 and Sox2 in GABA-ergic neurons. The majority of cell types have a high Pearson correlation (>0.8) between matched cell types’ average expression profile; even for the rare cell-type microglia, the correlation remains reasonably high (0.75) (**Fig. 1d**). We are also able to distinguish early maturing oligodendrocytes in the seqFISH data based on Itpr2 expression (**Fig. 1c**, OPC column) as previously reported by Zeisel et al^15^. Inhibitory GABA-ergic neurons and excitatory glutamatergic neurons exhibit strong anti-correlation to each other (**Fig. 1d**).

As an additional validation, we examined the Nissl and DAPI staining images which are known to have distinct patterns between astrocytes and neuronal cell types. As Nissl is a neuronal stain and DAPI stains DNA, astrocytes are typically associated with DAPI but not Nissl, whereas neurons are stained for both. Our cell-type mapping results highly agree with these patterns. Over 89% of predicted astrocytes exhibit strong DAPI staining but weak or no Nissl staining across cortex columns (**Supplementary Notes**, **Supplementary Table 3**). Taken together, these analyses indicate that the majority of cells were mapped to the correct cell types. By combining cell type predictions from scRNAseq and positional information from seqFISH, we were able to construct a single-cell resolution landscape of cell type spatial distribution (**Fig. 1e**). As expected, this landscape is very complex, with different cell types intermixed with each other (**Fig. 1e**). On the other hand, it is clear that there remains significant heterogeneity within each cell-type.

### A systematic approach to identify multicellular niche from spatial genomics data

Microenvironment in tissues can contribute to heterogeneity in addition to cell type specific expression patterns. To systematically dissect the contributions of microenvironments on gene expression variation, we developed a novel hidden-Markov random field (HMRF) approach^32^ to unbiasedly inform the organizational structure of the visual cortex. An overview of this approach is illustrated in **Fig. 2a**. The basic assumption is that the visual cortex can be divided into domains with coherent gene expression patterns. A domain may be formed by a cluster of cells from the same cell-type, but it may also consist of multiple cell-types. In the latter scenario, the expression patterns of cell-type specific genes may not be spatially coherent, but environment-associated genes would express in spatial domains. HMRF enables the detection of spatial domains by systematically comparing the gene signature of each cell with its surroundings to search for coherent patterns. Briefly, we computationally constructed an undirected graph to represent the spatial relationship among the cells, connecting any pair of cells that are immediate neighbors (**Fig 2a, b**). Each cell is represented as a node in this graph. The domain state of each cell is influenced by two sources (**Fig 2b**): 1) its gene expression pattern, and 2) the domain states of neighboring cells. The total contribution of neighboring cells can be mathematically represented as a continuous energy field, and the optimal solution is identified by searching for the equilibrium of the field (see Methods, Supplementary Note X for mathematical details). Next we applied our HMRF model to analyze the 1597-cell mouse visual cortex seqFISH dataset. The expression of the 125 genes ranges from being highly scattered to spatially organized. To enhance spatial domain detection, we defined a spatial coherence score, and selected the top 80 genes for HMRF analysis (see Methods). As an additional filter, we further removed 11 genes that are highly specific to a single cell type, resulting 69 genes (**Supplementary Table 4**) for spatial domain identification. We found this additional filtering step improves the resolution while preserving the overall spatial pattern (**Supplementary Fig 2**).

**Figure 2.**
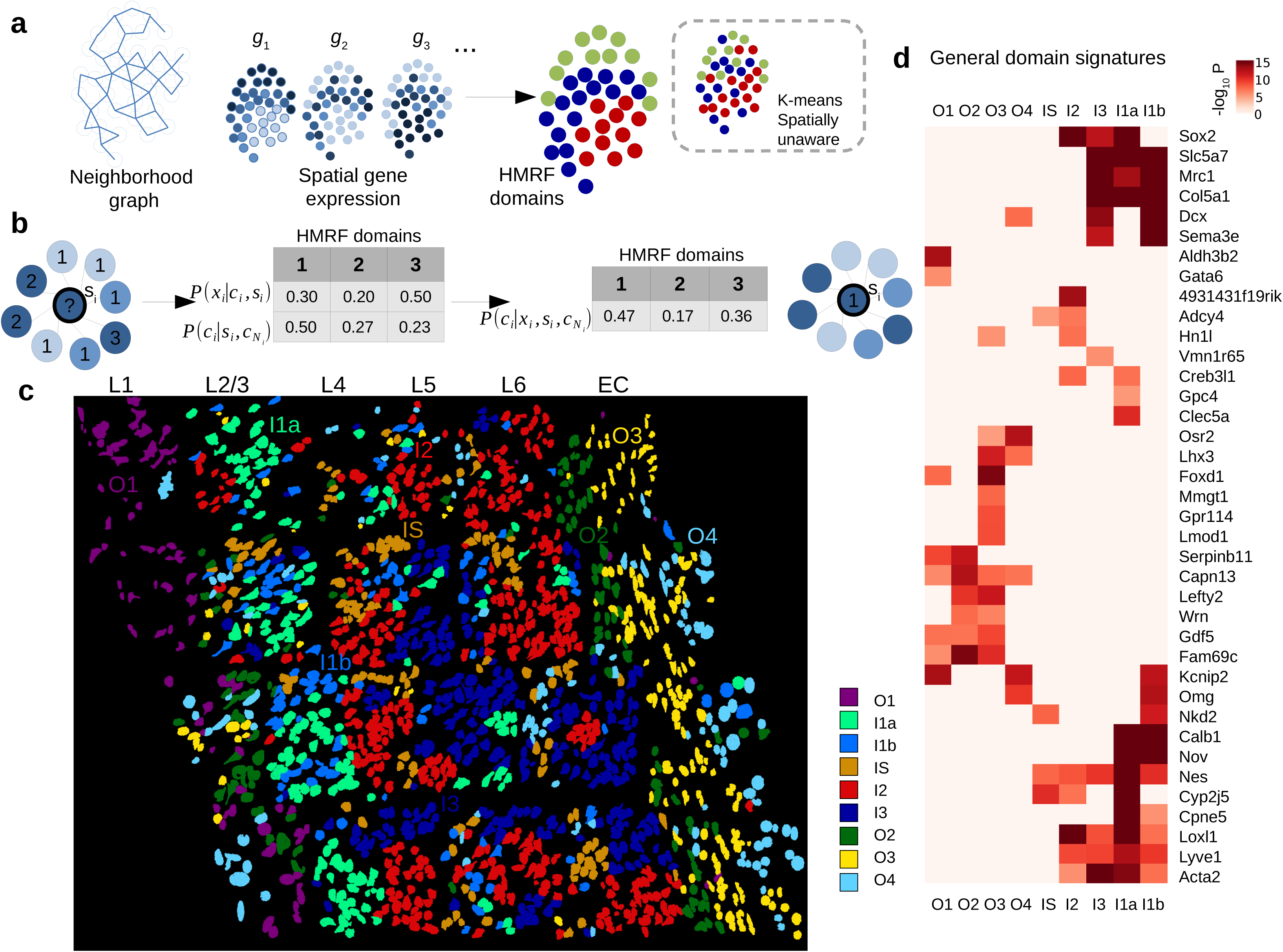
Spatial domain dissection in seqFISH data using hidden Markov random field (HMRF) approach. a. A schematic overview of the HMRF model. A neighborhood graph represents the spatial relationship between imaged cells (indicated by the circles) in the seqFISH data. The edges connect cells that are neighboring to each other. seqFISH-detected multigene expression profiles are used together with the graph topology to identify spatial domains. In contrast, k-means and other clustering methods do not utilize spatial information and therefore the results are expected to be less coherent (illustrated in the dashed box). b. An intuitive illustration of the basic principles in a HMRF model. For a hypothetical cell (indicated by the question mark), its spatial domain assignment is inferred from combining information from gene expression (x_i_) and neighborhood configuration (c_Ni_). The color of each node represents cell’s expression and the number inside each node is domain number. In this hypothetical example, combining such information results the cell being assigned to domain 1, instead of domain 3 (see Methods). c. HMRF identifies spatial domain configuration in the mouse visual cortex region. Distinct domains reveal a resemblance to layer organization of cortex. Naming of domains: I1a, I1b, I2, I3 are inner domains distributed in the inner layers. O1-O4 are outer domains. IS is inner scattered state. These domains are associated with cell morphological features such as distinct cell shape differences in outer layer domains. Cell shape information is obtained from segmentation of cells from images (see Methods). d. General domain signatures that are shared between cells within domains.

HMRF modeling of the visual cortex region revealed 9 spatial domains (**Fig. 2c**). These domains have distinct spatial patterns; some display a layered organization that resembles the anatomical structure^33^. For example, four of the domains are located on the outer layers of the cortex therefore labeled as O1, O2, O3, and O4, respectively (**Fig. 2c**). The locations of these layers roughly correspond to the well-characterized L1, L6, and external capsule (EC) layers, respectively. Four domains are located on the inside of the cortex therefore labeled as I1a, I1b, I2, and I3, respectively (**Fig. 2c**). These domains roughly correspond to the L2-5 layers. These inner domains are less pronounced than the outer domains, which is consistent with previous anatomical analysis. Finally, one domain is sporadically distributed across in the inner layers of the cortex, therefore labeled as IS (**Fig 2c**). Of note, such domain-like patterns are not visible in the cell-type localization pattern (Fig 1e). Consistent with these results, t-SNE plot using these 69 genes identified clustering patterns similar to the domain annotations but differ greatly from the cell-type annotations (**Supplementary Fig. 3**). These results strongly suggest HMRF provides complementary information to cell type annotations.

By overlaying cell type annotations, we see that each domain generally consists of a mixture of GABA-ergic, glutamatergic neurons and astrocytes interacting in each environment (e.g. domain I1a in **Supplementary Fig. 4**). The decomposition of mouse visual cortex into spatial domains suggests that a spatial gene expression program is shared across cells in proximity. Differential gene expression analysis identified distinct signatures, which we term as the general domain signatures, associated with each spatial domain (**Fig. 2d, Supplementary Figs. 5, 6, 7**). For example, genes Calb1, Cpne5, Nov are preferentially expressed in inner domains (I1a, I1b), whereas genes Serpinb11, Capn13 are highly enriched in outer domains (O1, O2). Different outer domains can be further distinguished by additional markers, such as Mmgt1 (O3), Aldh3b2 (O1), and Fam69c (O2). Importantly, these spatial gene signatures transcend multiple cell types therefore are distinct from cell-type specific signatures (**Supplementary Figs 6, 7**). The spatial marker genes are highly consistent with their spatial expression in Allen Brain Atlas^33^ ISH images, such as Calb1, Cpne5, Nov (see **Supplementary Fig. 8**). Other markers such as Nell1, Aldh3b2, Gdf5 have layer-specific expressions that are consistent with Zeisel et al^15^ (**Supplementary Fig. 8**). We summarized the gene signature of each domain as a metagene, defined as the average expression of the subset of genes that are specifically associated with the domain. This provides an “analog” representation of the spatial domain information as an additional diagnostic (**Supplementary Fig. 9**). Taken together, these analyses strongly suggest that our model for analyzing seqFISH data is able to detect functionally and transcriptionally distinct spatial environments.

### Integrative analysis identified cell-type, environmental interactions

Glutamatergic neurons mediate the neuronal circuit in the visual cortex by playing a primarily excitatory function. It is also well-known that the behavior of different glutamatergic neurons can be very different^27, 34^. By combining cell-type mapping and spatial domain identification, we set out to dissect the source of heterogeneity within glutamatergic cells. First, nearly all glutamatergic cells express cell-type specific markers such as Gda and Tbr1 (**Fig 3a top**). In addition to demonstrating cell type identity, there exists substantial heterogeneity within glutamatergic cells in a spatially dependent manner. As glutamatergic cells are spread across all 9 domains, each subset expresses a different gene signature in accordance to domain annotation (**Fig. 3a middle, bottom**). First, the general domain signatures in Fig 2d, aggregated as metagenes, can separate glutamatergic cells into domains (**Fig 3a middle**). Secondly, beyond the general signature, an additional set of gene signatures are differentially expressed between glutamatergic cells in different domains (**Fig. 3a bottom**). To distinguish these genes from the general domain signatures which are cell-type transcending, we refer to these genes as the glutamatergic restricted signatures. For example, Mmp8 expression is restricted to domain O2 (**Fig 3a bottom**), whereas Hoxb8 expression is specific to O3, and Nfkb2 to IS (**Fig 3a bottom**). Collectively, the domain-specific signatures map out the spatial patterns of expression within glutamatergic cells, demonstrating their power to differentiate subgroups of this neuron class (**Supplementary Figs. 9, 10, 11**).

**Figure 3:**
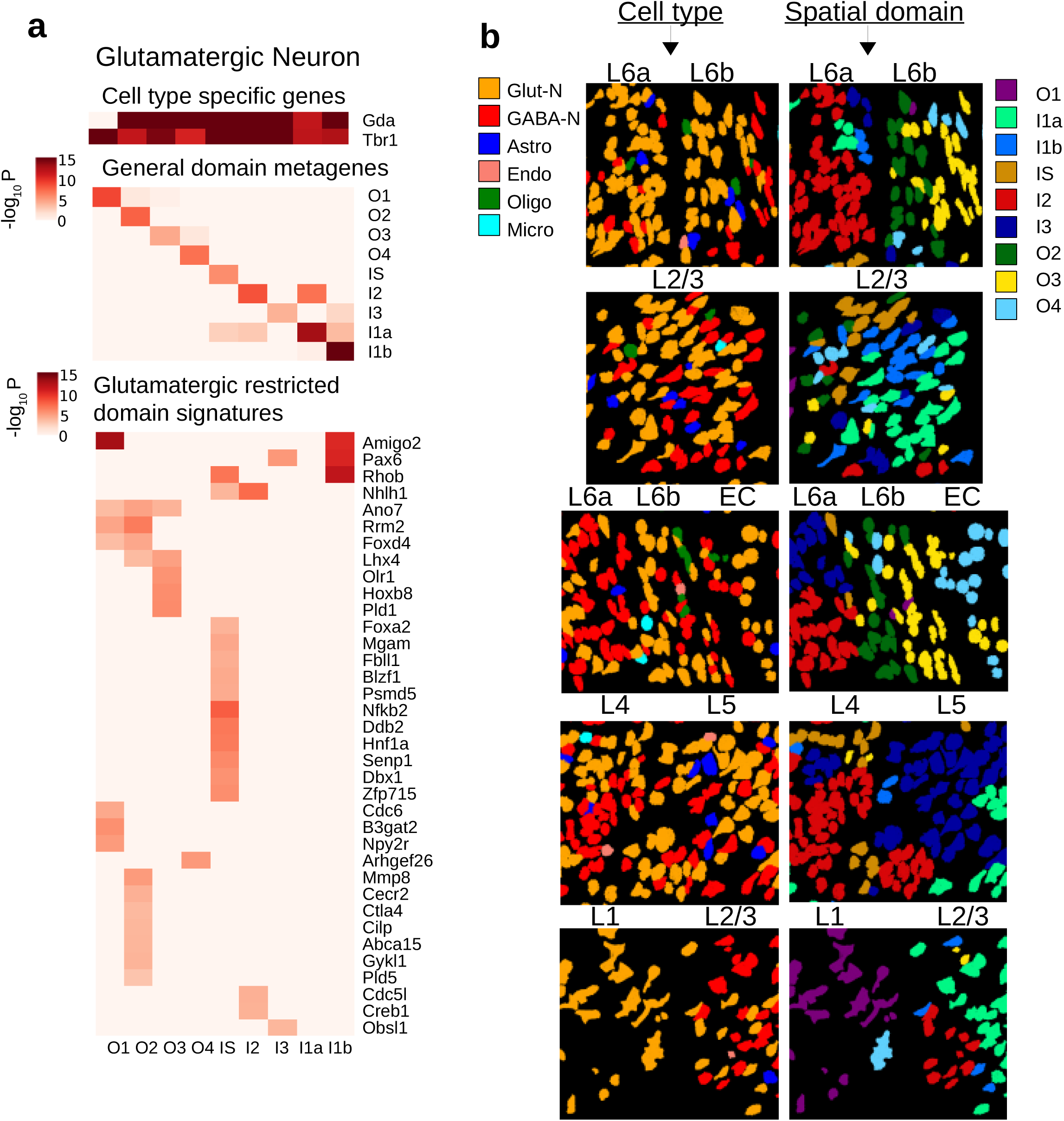
HMRF analysis identified domain associated heterogeneity within glutamatergic cells. a. Three major sources of variations in glutamatergic neurons. (Top): cell type specific signals Gda and Tbr1. (Middle): general domain signatures as in Fig 2d, summarized into metagenes’ expression. (Bottom): glutamatergic restricted domain signatures, found by comparing glutamatergic cells across domains and removing signatures that are general domain signatures. b. Snapshots of single cells. Each row is a snapshot of cells at the boundary of two layers. Each of two columns is a type of annotation: (left column) cell type, (right column) HMRF domains. Cell type is incapable of explaining layer-to-layer morphological variations: e.g. glutamatergic cells (orange) is present in all layers yet morphological differences exist within glutamatergic cells. HMRF domains better capture the boundary of two layers in each case, in that the domains can separate distinct morphologies

By visual inspection, we observed remarkable morphological switches near the boundary between different domains at multiple regions (the three groups of cells in panel L6a, L6b, EC of **Fig 3b**), including change of circularity and cell orientations, and accompanied by metagene expression switches (**Supplementary Fig 11**). To systematically compare the morphological differences between different domains, we extracted quantitative information of 15 different morphological features per cell based on the Nissl staining images, and compared the statistical distributions across different domains. Indeed, we found a number of features display strong domain associations, including circularity in O4 (P<6.1e-12), width in I1b (P<1.6e-14), angle in O3 (P<6.7e-18), minimum feret diameter in I1a (P<3.0e-11) (**Supplementary Fig 12**). Of note, these differences cannot be identified by using cell-type mapping alone (**Fig 3b**). Thus, within neuronal cell types, such as glutamatergic or GABA-ergic neurons, there remains significant morphological differences across domains, suggesting that spatial positions accounts for a large part of morphologies in these cells, consistent with known morphological diversity in the cortex. Overall, these analyses strongly suggest that spatial domain variation plays an important role in mediating cellular heterogeneity.

### Using HMRF domain information to reanalyze scRNAseq data

ScRNAseq data does not contain spatial information. However, using domain signatures derived from seqFISH as a guide, we were able to infer spatial locations from scRNAseq data. In order to dissect the contribution of environmental factors to transcriptomic heterogeneity, we focused on glutamatergic cells, and combined the general domain signatures with the additional set of markers that are glutamatergic restrictive. Using these expanded domain signatures (**Supplementary Table 5**) summarized as metagenes, we were able to uncover a hidden structure within the glutamatergic cells (**Fig 4a, b**). Strikingly, the glutamatergic cells can be partitioned into nine different clusters based on the expanded domain signatures, which were highly consistent with seqFISH data analysis (**Fig 4a, b**). As such, these clusters were labeled according to their enriched metagene signatures (**Fig 4a**).

**Figure 4.**
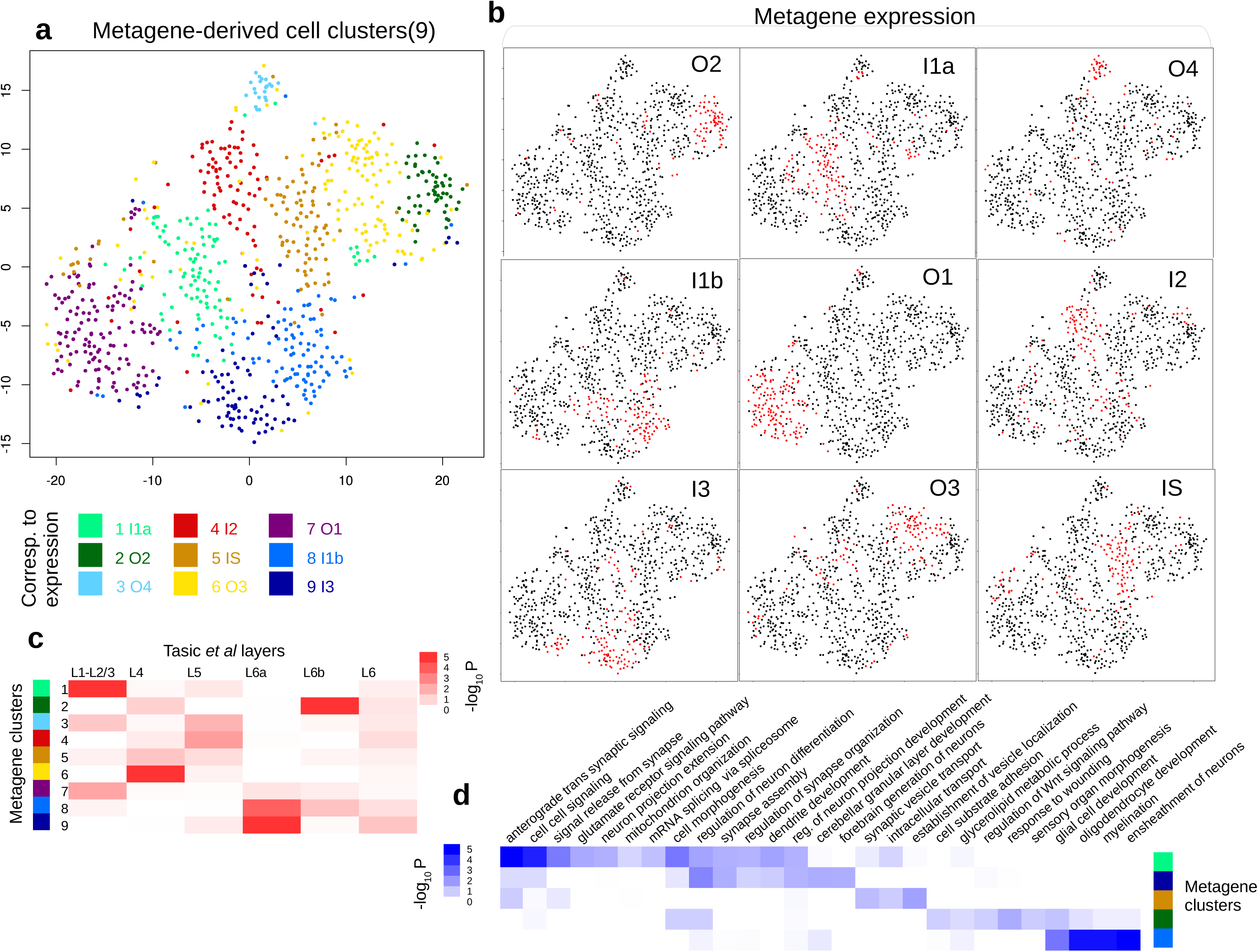
Reanalysis of single-cell RNAseq data (from Tasic *et al*) with domain signatures summarized into metagenes. a. t-SNE plot shows how glutamatergic cells from Tasic et al cluster according to expanded domain signatures aggregated as metagenes (shown in (b)). Colors indicate k-means clusters (k=9). Each cluster is annotated by its enriched metagene activity. b. Binarized metagene expression profiles for the glutamatergic cells. Red: population that highly expresses the metagene. c. Spatial clusters defined according metagenes are enriched in manual layer dissection annotations. Column: layer information obtained from microdissection. Row: metagene based cell clusters. d. Inferred spatial clusters of glutamatergic neurons are enriched in distinct GO biological processes.

We compared the inferred domain annotations with the original sites of dissection in Tasic et al. Several domains match the corresponding layer structure very well (**Fig 4c**). For example, cluster 1 (annotated as domain I1a based on metagene analysis) significantly overlaps with L1-L2/3 (P<2.3e-6). Cluster 2 (annotated as domain O2) overlaps with L6b (P<4.8e-9), and cluster 9 (annotated as domain I3) significantly overlaps with L6a dissection label (P<1.0e-8). On the other hand, clusters 3, 4, and 5 (annotated as domains O4, I2, and IS) do not correspond to specific layers.

Using the whole transcriptomes from scRNAseq, we searched for additional domain specific gene signature based on co-expression analysis. Our analysis identified a number of genes that were not assayed by seqFISH, including Tubb2a (I1a), Ndrg3 (O4). We examined the corresponding ISH images in the Allen Brain Atlas, and found that the inferred spatial patterns agree well with the imaging data (**Supplementary Fig. 13**). We further conducted gene set enrichment analysis based on the inferred domain-specific markers, and identified a number of functional biological processes that are enriched in specific domains (**Fig 4d**).

An important question is whether the distinction between the subpopulations identified through our integrative analysis simply reflects subtype differences which can be identified through scRNAseq analysis alone. To address this question, we systematically compared the domain and subtype annotations using a number of approaches, including the underlying gene signatures, the grouping of cells based on domain or cell subtype annotations, and tSNE-based visualizations (**Supplementary Figs 14,15**). Based on these comparison, our conclusion is two-fold. On on hand, we observed a non-negligible association between the two sets of annotations, such as at L6b_Serpinb11, L2/3_Ptgs2, L6a_Sla (**Supplementary Fig 14**). For example, several domain-specific markers are also markers of specific cell subtypes, such as Serpinb11, Cpne5, and Sema3e (**Supplementary Fig 16a**). On the other hand, it is also clear that the overall structure of domain-and subtype-annotations are very different. For example, cells inferred to be located in domains O1, IS, O4 spread across multiple subtypes (**Supplementary Figs 14, 16b**). Conversely, neither L5a_Batf3 nor L5a_Hsd11b1 subtype is associated with any specific domain (**Supplementary Fig 14**). Taken together, these analyses strongly indicated the domain patterns are distinct from, and therefore complementary to, cell subtype annotations. Thus, integrating seqFISH data analysis provides new insights into scRNAseq data.

### HMRF analysis reveals region-specific variation among astrocytes

Next, we investigated the environment effect on astrocytes, which are also known to have substantial heterogeneity^20, 35^. Our cell type mapping identified 47 astrocytes in the seqFISH data. These cells all expressed key astrocyte markers (**Fig 5, box 1**) but were spread across 5 different spatial domains (O1, O2, O3, I1a, and I3) (**Fig. 5**). Of note, a number of astrocyte markers^20^ (**Supplementary Fig. 17**) are only expressed in specific domains (**Fig. 5**). As an example, Acta2, Col5a1, and Sox2 are strongly associated with domain I1a, while their expression levels are greatly reduced in domains O1 and O2. On the other hand, the expression levels of Clec5a and Ankle1 are high in domains O2 and O1 but much lower in other domains. The spatially dependent variations may underline important functional differences.

**Figure 5:**
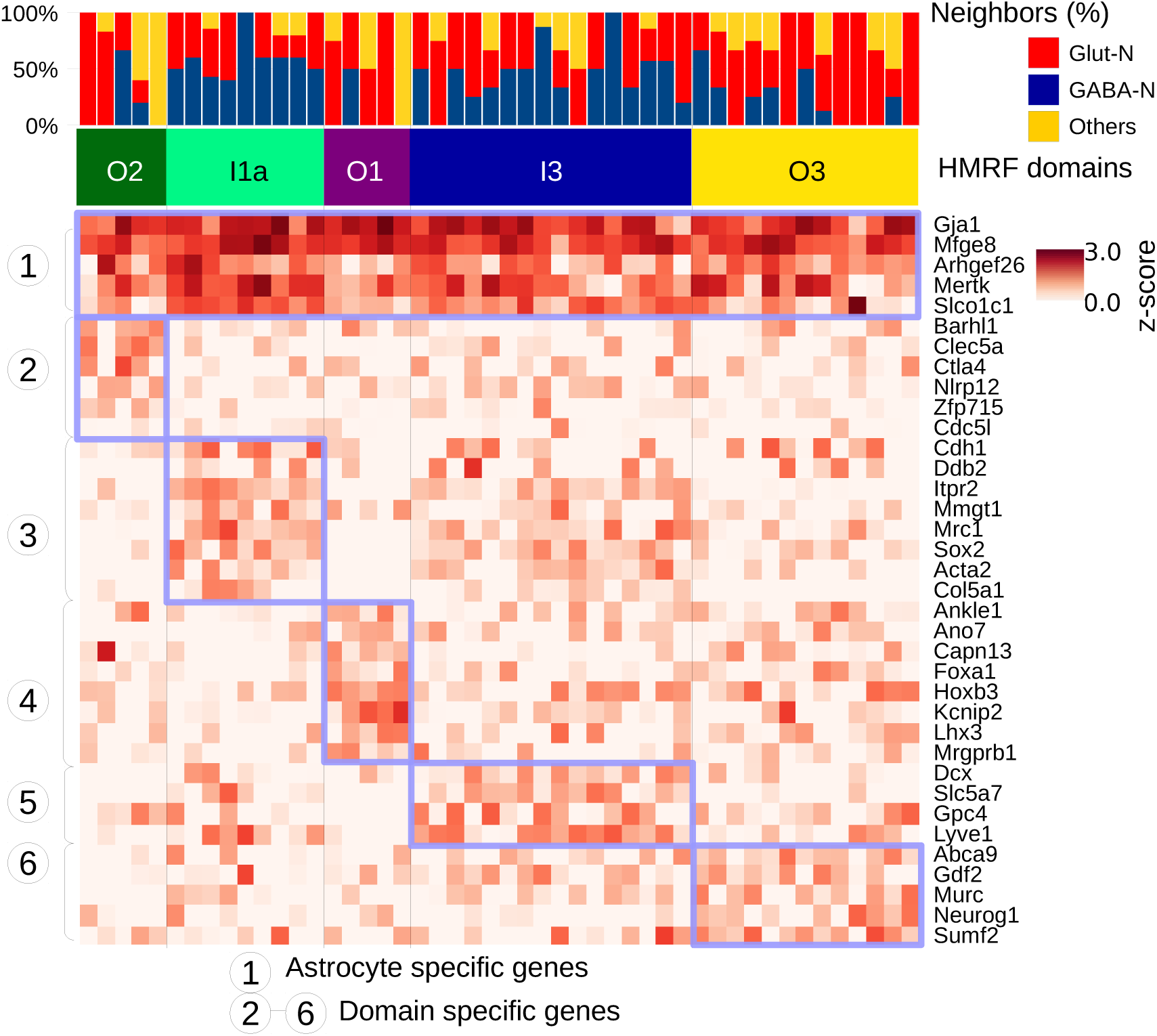
Spatially dependent astrocyte variation revealed by HMRF. Neighborhood cell type composition for the 47 astrocyte cells (columns). Cells are ordered by HMRF domain annotations. The heatmap shows single cell expression of astrocytes clustered by domain-specific genes. Blue-box highlights the common signatures expressed in each domain’s astrocyte population.

## Conclusion

A major goal in single-cell analysis is to systematically dissect the contributions of cell-types and environment on mediating cell-state variability. To achieve this goal, we presented an HMRF-based computational approach to combine the strengths of sequencing and imaging-based single-cell transcriptomic profiling strategies. We showed that our method can be used to correctly detect spatial domains in the mouse visual cortex region. In doing so, we were able to identify environment-associated variations within a common cell-type. Our analysis also demonstrated that novel insights can be gleaned from single-cell data by an integration of information from complementary technologies. In particular, integrating scRNAseq data allows us to map cell-types more accurately than in seqFISH data analysis, whereas integrating seqFISH data allows us to extract spatial structure in scRNAseq data analysis. To test the generalizability of our method, we applied it to analyze a published spatial transcriptomic dataset obtained from a very different technology^36^. Here, spatial information was identified by hybridizing mRNA to a specially designed tissue-microarray containing spatial barcoding oligo-probes. Despite the significant platform differences, our HMRF model was able to recapitulate the spatial domains that are consistent with the underlying anatomical structures (**Supplementary Fig 18**). In another example, we analyzed seqFISH data^19^ obtained from a different region (dentate gyrus) using different probes. Again the results are consistent with the anatomical structure (**Supplementary Fig 19**). These analyses strongly indicate our method is generally applicable. Future work will continue to investigate the mechanisms underlying the interactions between cell-type and microenvironment.

## Author Contributions

Conception and supervision of project: G.C.Y., L.C. Conception of HMRF and SVM models: Q.Z., G.C.Y. Conducting and supervision of computational analyses: Q.Z, G.C.Y. Conducting and supervision of seqFISH experiments: S.S, L.C. Writing: Q.Z., S.S., R.D., G.C.Y., L.C. All authors contributed ideas for this work. All authors reviewed and approved the manuscript. This research was supported by a Claudia Barr Award, a Chan Zuckerberg Initiative Award, and NIH grant R01HL119099 to G.C.Y. and NIH R01 HD075605 to L.C.

## Methods

### SeqFISH data generation

SeqFISH data in the mouse visual cortex region was generated as described previously (Shah 2016). Briefly, 100 genes were encoded using a temporal barcoding method and 25 genes were quantified individually. To encode 100 genes, 4 rounds of hybridization were performed using 5 distinct fluorescence channels. Out of a total possible 625 barcodes, 100 were chosen such that loss of signal in any given hybridization still allows accurate decoding of the spot. Every transcript was hybridized in every round using a given probe set. After hybridization, the signal was amplified using smHCR and images were taken at predefined locations in the mouse visual cortex. The DNA probes along with the amplification polymers were digested using DNase I DNAseI leaving behind a naked RNA for re-hybridization with the next probe set. A round of imaging with DAPI staining (which labels the DNA) was done before any RNA hybridization to image all nuclei in the fields and a final round of Nissl staining (which labels the cell body in neuronal cells) was imaged to identify cell boundaries. Cells were segmented based on DAPI staining, Nissl staining, and RNA point density. Once all imaging rounds were completed, these images were aligned using a 2D normalized cross correlation and each spot was decoded based on the unique color switching pattern. For the 25 genes that were labelled without any encoding, simple spot counting was done to identify the number of transcripts. These transcripts were then assigned to cells based on the location of the transcript and the segmentation masks. For more details regarding the seqFISH method, please refer to Shah et al. 2016^19^. The spatial coordinates of the cells are provided in **Supplementary Data**.

### SeqFISH data normalization and bias correction

The seqFISH gene expression matrix, represented by log(count + 1), was normalized by row and column z-scoring to remove cell-specific and gene-specific biases. Potential field imaging biases were estimated and removed by using a multi-image regression algorithm similar as previously done^37^. Briefly, for each gene, the imaging bias at each binned location was estimated by averaging the normalized gene expression levels over 8 neighboring bins within each field followed by averaging across all fields. The estimated bias was then modeled by principal component analysis (PCA). The contributions of the four most significant PCs were estimated by linear regression and removed from the normalized gene expression matrix (**Supplementary Fig 20**).

### Cell type mapping

Single-cell RNAseq data for the mouse visual cortex were obtained from Gene Expression Omnibus^38^ (GSE71585). Cell-type information corresponding to 1723 cells was obtained from the original paper^27^ (Tasic 2016). In this analysis, we considered the 8 major cell types: GABAergic, glutamatergic, astrocytes, 3 oligodendrocyte groups, microglia, and endothelial cells. Differentially expressed genes among different cell types were identified by MAST^39^. We trained classifiers of cell types from single-cell RNAseq dataset by using the multiclass SVM formulation. For each cell-type, we built a classifier as follows. Let *x*_*i*_ *, i*= 1, …, *n*, be the gene expression pattern for the *i*-th cell, and *y*_*i*_ code for cell-type identity: *y*_*i*_ =1 if cell i belongs to the specified cell type and −1 otherwise. We selected the linear kernel that produces two hyperplanes that best separates the two classes. The objective function is defined as follows

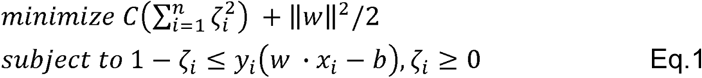

Here *w* is the normal vector to the hyperplane used to represent margin. The squared hinge loss function 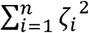 is used here to quantify the margin of misclassification error. *C* is a regularization parameter that trades off misclassification due to overfitting against simplicity of the decision function. A lower *C* increases the ability of the model to generalize to unseen data at a cost of larger fitting error. For testing data, the sign of *w − x*_*i*_ *-b* is used to predict cell type identity. We used the Python LinearSVC implementation, which is part of the scikit-learn 0.19 library^40^, with the following parameter setting: class_weights=balanced, dual=False, max_iter=10000, and tol=1e-4.

Using the SVM model formulated as above, we first tested how many genes are needed for accurate cell-mapping. To this end, we randomly subset 20, 40, 60, and 80 genes from the list of differentially expressed genes and, for each gene set, built a vanilla SVM classification model to map each cell in the single-cell RNAseq dataset to its corresponding cell-type. The cross-validation accuracy was evaluated by using 4-fold cross-validation. Our results indicated that a high accuracy (>90%) can be obtained with 40 or more genes.

In addition to the major cell types mentioned above, Tasic et al also identified 22 fine cell classes, and 49 minor cell classes. Using the same approach, we also evaluated the accuracy of refined cell-type mapping (**Supplementary Fig 21**). We found that approximately 200 genes are required to achieve 85% accuracy in predicting 22 finer classes, and over 800 genes are needed to predict the 49 minor cell types with 75% accuracy. Therefore, we focused on the mapping of 8 major cell types on seqFISH given that they can be predicted accurately with fewer than 100 genes (ROC curves in **Supplementary Fig 22**).

To map cell-types in the seqFISH data, we made a few modifications to incorporate the platform differences. First, since 125 genes were profiled by seqFISH, we used the top differentially expressed genes (p<1e-20) in the scRNAseq dataset for cell-type mapping. Based on the subsampling analysis described above, these 43 genes were sufficient for accurate cell-type mapping. Second, the scRNAseq data were z-score transformed so that the dynamic range was comparable with seqFISH data. Third, we used quantile normalization^30^ to convert seqFISH data so that the statistical distribution was almost identical to single-cell RNAseq data.Fourth, we chose the regularization parameter *C* to maximize the cross-platform correlation between the cell-type specific gene expression profiles, resulting an estimate of *C*=1e-6. Finally, to account for the possibility that certain cells cannot be unequivocally assigned to a single cell-type, we used Platt scaling^31^ to convert SVM output to a probability measure and then selected a cutoff value of 0.5 probability to filter cells that can be confidently mapped to a single cell-type. 97 (5%) cells did not pass this filter.

### Hidden Markov random field

Hidden Markov random field (HMRF) is a graph-based model commonly used for pattern recognition in image data analyses^32, 41^. In a common setting, HMRF is used to model the spatial distribution of a signal, such as the pixel intensities over a 2D image. The spatial structure is represented as a set of nodes on a regular grid, where neighboring nodes are connected to each other. The spatial pattern is “hidden” in the sense that it must be indirectly estimated from other variables that can be directly measured. The most important assumption in HMRF is the Markov property, which states that the spatial constraints can be reduced to considering only correlation between immediate neighboring nodes. This simplifying assumption implies that the joint distribution can be decomposed as products of much smaller components each defined on a fully connected subgraph (termed cliques). As has been done previously, we decomposed the graph into size-2 components (or edges in the graph) that provides a convenient means to estimating the MRF by using pairwise energies.

Specifically, let *S=* {*s*_*i*_} be the nodes in the graph. The set of nodes and the adjacency relation as defined by the local neighborhood graph forms the neighborhood system (*S*, {*N*_*i*_}). Every node is associated with observed signal values *x*_*i*_. Let *c* = {*c_i_ =1,…, K*} represent the set of possible classes of patterns. The joint probability that a node si is associated with class is specified by the following equation:

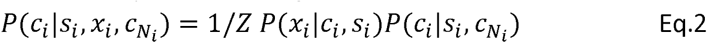

In the right hand side, the term *P*(*x*_*i*_ *|c*_*i*_ *, s*_*i*_) models the effect of the node *s_i_*’s own gene expression, whereas 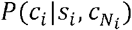 models the effect of the neighboring cells configuration 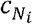. The combined effect of these two terms is schematically shown in **Fig. 2**. The latter term is further determined by the Gibbs distribution:

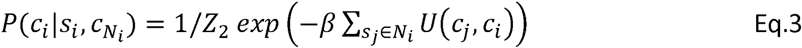

where *U*(*c*_*j*_ *, c*_*i*_) is referred to as the energy function. The exact formulation of *U*(*c*_*j*_ *, c*_*i*_) is dependent on the specific application, and it imposes the assumption of how neighboring nodes are interacting with each other. Here we use the special case Pott’s model.

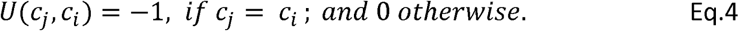

which means that the effects of neighboring cells are additive. Essentially, 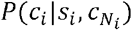 expresses the total energies as a summation of pairwise interaction energies with neighbors. The parameter beta reflects the strength of interactions.

### Application to seqFISH data

The HMRF model described above is naturally applicable to analyze seqFISH data. Here each class of patterns corresponds to a spatial domain. The observed signals are gene expression levels measured by seqFISH data, whose distribution is modeled as a multivariate Gaussian random variable. The application of HMRF to seqFISH data analysis involves the following 4 components. 1) Neighboring graph representation. 2) Gene selection. 3) Domain number selection, and 4) Implementation and model inference. The details of each component are described below.

1. Neighborhood graph representation. An undirected graph was constructed to represent the spatial relationship between the cells. Each node represents a cell, and each edge connects a pair of neighboring cells. The neighborhood size was chosen such as on average each cell has five neighboring cells.
2. Gene selection. We selected a subset of genes whose expression patterns tend to be spatially coherent based on the following analysis. For each gene *g*, cells were divided into two mutually exclusive sets: the first set, denoted by L1, contains cells with high expression at the 90th percentile expression level cutoff, and the rest of the cells were denoted by L0. The spatial coherence of gene expression was quantified as the Silhouette coefficient^42^ of the spatial distance associated with these two cell sets. Specifically, the Silhouette coefficient is calculated as:

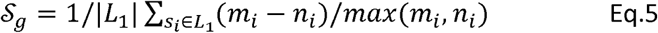

where for a given cell *S_i_* in Set L1, *m*_*i*_ is defined as the average distance between *s*_*i*_ and any cell in L0, and *n*_*i*_ is defined as the average distance between and any other cell in L1. Here, we used the rank-normalized, exponentially transformed distance to quantify the local physical distance between two cells. For a pair of cells si and sj, this distance is defined as 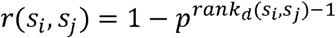 where is the mutual rank^43^ of *s*_*i*_ and *s*_*j*_ distances {Euc(si,*)} and {Euc(sj,*)}. Hence, this exponentially weighted function^44^ is designed to give more emphasis on closely located cells and penalizing far-away cells’ distance to a large number. p is a rank-weighting constant (0<p<1.0) set at 0.95. The statistical significance of 𝒮 _g_ was evaluated by random permutation, and the genes associated with significant values of 𝒮 _g_(p-value < 0.05) were selected as spatially coherent. Using the above criteria, we found 80 spatially coherent genes. We further removed 11 cell type specific genes (MAST P<1e-20) which have average expression z-score >2. We found this additional filtering step is useful for improving the resolution while preserving the overall spatial pattern (**Supplementary Fig 2**). We repeated the analysis using varying degree of stringency for selecting spatially coherent genes (**Supplementary Fig 23**), varying the degree of excluding cell-type specific genes (**Supplementary Fig 2**), and varying beta (**Supplementary Fig 24**), and found that the overall patterns identified by the HMRF model is robust against these variations.
3. Domain number selection. We used k-means clustering results as initialization for the HMRF domains. The value of k was selected based on the gap-statistics^45^.
4. Implementation and model inference. The model parameters were inferred by using the Expectation-Maximization (EM) algorithm^46^. We developed a new implementation based on the MRITC R package^47^ and GraphColoring Java package^48^. The implementation contains modifications to accommodate arbitrary neighborhood graph topology. The domain assignment for each cell was determined by using *maximum a posteriori* estimation, which can be viewed as the equilibrium state of the energy function. See **Supplementary Notes** for implementation details.

## Robustness analysis of the HMRF model

We also tested the robustness of our HMRF model against spatial perturbation. This was achieved by randomly exchanging the spatial locations of a subset of cells (10%, 20%, 40%, 100%). At 100% exchanging rate, the spatial coherence is completely disrupted. Log-likelihood of the HMRF model was recorded and compared across scenarios. As expected, the log-likelihood achieves maximum at a low perturbation rate and gradually decreases as the exchange rate increases. The difference between the perturbed and unperturbed data is highly statistically significant (**Supplementary Fig 25**).

## Domain-specific gene signatures

For each spatial domain, we identified a subset of genes that were significantly up-regulated in the domain compared to cells in other regions. Specifically, we require that the selected gene be both significant in one-vs-one tests (comparing it to one domain at a time, and pass significance threshold P<0.05 in at least 7 of 8 such tests, Welch’s t-test) and significant in one-vs-rest test (P<1e-5 Welch’s t-test). The use of t-test is justified as the expression z-scores are normally distributed (**Supplementary Fig 26**). Non-parametric Mann-Whitney U tests yield similar signatures (**Supplementary Fig 27**). Accordingly, we defined a metagene signature as the average expression level for this subset of up-regulated genes. These domain-associated metagene signatures (as appears in **Fig 2d**) transcend cell types (**Supplementary Figs 6,7**). Furthermore, we restricted this comparison to each specific cell type, and obtained an additional list of genes that are differentially expressed between domains. An expanded domain-metagene signatures was then defined based on the merged gene subsets. For glutamatergic cells, the expanded metagene signatures are summarized in **Supplementary Table 5**.

## Analysis of spatial structure in the single-cell RNAseq data

In order to systematically characterize the spatial structure within a scRNAseq data, we summarized the gene signature associated with each spatial domain as a metagene (as described in the previous section). For simplicity, the overall expression of an expanded domain-specific metagene signature in each cell was quantified as the mean z-scored expression of all constituent genes in the signature. A t-SNE analysis was performed on this matrix using the Rtsne package with parameters pca_scale=T, perplexity=35. Cell subpopulations with similar metagene expression patterns were identified by K-means clustering analysis (K=9). We next annotated each cluster as belonging to the expression of one metagene. By comparing the binarized metagene expression population (**Fig 4b**) and the K-means cluster annotations (**Fig 4a**), all of the K-means clusters were assigned as uniquely belonging to one metagene.

For each subpopulation discovered from metagene clustering above, we found differentially expressed (DE) genes for the population (2-sample t-test, unequal variance, P<0.05). With the DE genes, we carried out Gene Ontology enrichment analysis (using hypergeometric test) for each of 9 subpopulations to construct a functional enrichment profile in **Fig. 4** (hypergeometric test, top 500 DE genes analyzed per group, multiple hypothesis^49^ corrected P<0.05). Here we used genes expressed in glutamatergic cells as the background gene-set when doing enrichment analysis.

Tasic et al also provides layer information for a glutamatergic cell subset based on the layer from which the cells were manually dissected using different Cre-lines. To test whether the extracted subpopulation based on metagenes is enriched for a certain manually dissected layer of cells, we also performed hypergeometric test corrected for multiple hypothesis comparing manual annotations of cells to our HMRF domain-based annotations.

## Visualization of spatial domain and cell type specific variations

We created box plots to visualize the range of expression values for cells in different domains and for different cell types. Additionally, to see cell type transcending effect of domain signature genes, for each genes, we grouped cells by (cell type, spatial domain) pair, and plotted the expression distribution across groups ordered by spatial domains. Groups with less than 4 cells are removed as these skewed the comparison.

## Morphological analysis

We loaded the cell segmentations as regions of interest files (ROI) in ImageJ^50^, then used the Measure tool available in ImageJ to quantitatively measure over 15 morphological features for individual cells. We compared the distributions across different cell-types by using the Kolmogorov–Smirnov test. Statistical significance is judged by both 1) significance in at least 7 of 8 one-vs-one tests (P<0.05 per test), and 2) significance in one-vs-rest test (P<0.0001).

## Code Availability

Code is deposited at https://bitbucket.org/qzhudfci/smfishhmrf-py/.

## Data Availability

Expression data, spatial coordinates, SVM predictions, HMRF domains, and expression box-plots categorized by domains and cell types are deposited at https://spatial.rc.fas.harvard.edu.

